# Sexually dimorphic auditory representation in *Aedes aegypti* brains

**DOI:** 10.1101/2024.07.07.602439

**Authors:** Takuro S. Ohashi, Yifeng Y.J. Xu, Shunsuke Shigaki, Yukiko Nakamura, Tai-Ting Lee, YuMin M. Loh, Emi Mishiro-Sato, Daniel F. Eberl, Matthew P. Su, Azusa Kamikouchi

## Abstract

Male attraction to female flight sounds is a vital, reproducible component of courtship in many species of mosquitoes; however, female acoustic behaviours have proven challenging to define. To investigate sexual dimorphisms in acoustic behaviours, previous reports have largely focused on differences in mosquito peripheral ear anatomy and function. Whilst molecular investigations have recently begun on the auditory periphery, sexual dimorphisms in central processing of acoustic information have not yet been explored. Here we used a combination of neurotracing, calcium imaging and molecular analyses to examine sexual dimorphisms in auditory processing in the yellow fever mosquito *Aedes aegypti*. We identified shared and dimorphic neurons connecting male and female ears to the primary auditory processing centre in the brain, and defined multiple distinct neuronal clusters based on responses to auditory stimulation. We finally used transcriptomic and proteomic analyses to investigate the molecular factors underlying these differences, with motile ciliary-related terms significantly enriched in males.

## Introduction

Sexual dimorphisms in behavioural traits resulting from natural and sexual selection are found in many animals^1^, including differences in mating, feeding, aggression, and escape behaviours. These differences often rely on dimorphisms in sensory processing systems, consisting of peripheral sensory organs and central neural circuits^2–4^. In many animals, sensory organs have diverged to tune responses to important cues for each sex^5–11^. The importance of sexual dimorphisms in central sensory processing for sex-specific behaviours has also been reported^4,12–14^, though these insights are mainly limited to specific behaviours in certain model organisms.

Hearing systems frequently show sexual dimorphisms due to asymmetric mating behaviours. In torrent frogs, the male’s eardrum can sense a higher frequency of tones compared to females, allowing males to listen for female’s ultrasonic calls^15,16^. The yellow fever mosquito *Aedes aegypti* (*Ae*. *aegypti*), a major vector of various diseases^17,18^, also shows sexual dimorphisms in hearing behaviours, with male attraction to sounds (phonotaxis) playing a key role in mating behaviour. Mosquito mating largely occurs in swarms, male-dominated aggregations which females enter to copulate^19^. Males approach females by detecting their sexually dimorphic wing beat frequencies (WBFs), with males producing WBFs several hundred Hertz (Hz) higher than females^20,21^; male ears are tuned to this lower frequency female flight tone, facilitating phonotaxis. Female mosquitoes on the other hand do not appear to show attraction to male flight sounds and their hearing behaviours in general remain largely unclear^22,23^.

The mosquito hearing organ is located in the antennae^24^ and comprises two components: (i) the flagellum, a sound sail that vibrates with mechanical stimulation, and (ii) the Johnston’s organ (JO), the site of auditory transduction. The male flagellum is densely covered with fibrillae, whilst female have only sparse fibrillae distributions^25^. The mechanosensory neurons in JO (denoted JO neurons) are bipolar neurons, each having a single distal dendrite connected to the base of flagellum and an axon projecting to the brain^19,26,27^. Male *Ae. aegypti* JOs contain >15,000 JO neurons, making them the largest chordotonal organs reported in insects, whilst female JOs contain half the male number of neurons^26,28,29^.

In addition to anatomical differences between male and female ears, functional sexual dimorphisms also exist. Frequency matching between the female flight tone and the male peak flagellar vibration (referred to as mechanical tuning), both 400-500 Hz, facilitates male detection of the female’s flight tone. The peak oscillation frequency of the female flagellum on the other hand is ∼225 Hz^9,30,31^, potentially linked to predator detection but seemingly not to localization of conspecific males. Furthermore, male (but not female) mosquitoes can demonstrate large amplitude, seemingly mono-frequent flagellar oscillations referred to as self-sustained oscillations (SSOs). SSOs entrain only to pure tones close to female WBFs^9^, suggesting that SSOs in males play a key role in the acoustic processing of female flight tones.

Investigating this acoustic processing is made more complicated by differences in the peak sensitivity of electrical responses recorded from the axon bundle of male JO neurons (referred to as electrical tuning) to the aforementioned mechanical tuning of the flagellum^32,33^. This mismatch has led researchers to suggest that male JO neurons respond most sensitively not to female flight tones, but to non-linear phantom tones resulting from interactions between male and female WBFs, referred to as distortion products (DPs)^9,22,27^. Two prominent DPs, the quadratic distortion product (QDP; difference between male and female WBF) and the cubic distortion product (CDP; difference between twice the female WBF and the male WBF), are similar in frequency to the peak of the male electrical response^22,34^, supporting the hypothesis of a DP-based acoustic communication system^9,22^. Females have not been reported to show differences in mechanical and electrical tuning, though this is yet to be confirmed (but see Lapshin and Vorontsov 2013^35^).

Sexual dimorphisms at peripheral anatomical and functional levels are thus well explored, though the molecular bases of these differences have rarely been tested^36^. Investigations into central differences in mosquito auditory processing also have not yet been conducted. In fruit flies, axons of JO neurons innervate the antennal mechanosensory and motor center (AMMC) in the brain^37^. In mosquitoes, JO neurons have been reported to project to a part of the antennal lobe (AL) denoted as JO-C^38^; however, a recent study found *Ae. aegypti* JO neurons mainly project to a homologous region of the AMMC^29^. In several mosquito species, including *Ae. aegypti*, the size of the male AMMC is approximately double that of the female^39^, suggesting that the acoustic processing properties of the AMMC may also vary between sexes. Potential sexual differences in central processing, as well as the molecular basis of auditory processing in general, have yet to be elucidated however.

In this study, we sought to investigate central processing of auditory information in *Ae. aegypti* mosquitoes. We first compared the projection patterns of JO neuron axons in the brains of male and female *Ae. aegypti* and found that male JO neurons had more branches compared to females. Next, we compared the auditory response properties of the AMMC using calcium imaging, identifying sex-specific frequency properties and male-specific functional diversification. We also show that the low-frequency tones that elicit peak responses from female AMMCs can elicit negative responses in specific male neurons. Development of a quadratic state-space model suggested that males exhibiting SSOs can exhibit anti-resonance, dampening male flagellar vibrations at specific frequencies similar to those inhibiting specific male neurons.

To explore the molecular basis of observed sex differences, we also created transcriptomic and proteomic profiles of male and female ears, finding that cilium-related genes were highly overexpressed in male ears at both levels. Despite this, observations of the ciliary ultrastructure of JO neurons using electron microscopy found no obvious differences in dynein arm structures between the sexes. Our findings reveal different expression profiles of key molecules in mosquito hearing organs, and sexually dimorphic tonotopic maps in their AMMCs.

## Results

### Projection patterns of JO neurons in the brain of *Aedes aegypti*

The morphology of JO neurons in mosquitoes has been studied largely *via* fluorescent tracer injection and head sectioning^9,29,40^. However, examining the comprehensive projection pattern of JO neuron axons requires whole-brain imaging^37,41^. We therefore conducted whole-brain imaging using neural tracing from one side of the pedicel (JO neurons) to compare the panoptic morphology of JO neurons between sexes (see Methods). We also traced flagellar neurons to facilitate distinguishing projection patterns between different neuronal groups (flagellar and JO neurons in this case).

In both sexes, most of the fluorescent tracers from flagellum and a subset of tracers from pedicel labeled the antennal lobe (AL), known to be a primary olfactory processing center, matching previous studies (Supplementary Fig. 1)^38^. The tracer injected via the pedicel labelled axons in the AMMC (mainly from JO neurons) and some clusters of putative efferent neurons in the brain (Fig. 1)^40^. Males have at least three pairs (six) of putative efferent neuron clusters, while we found five such clusters in females (Fig. 1A, C). These clusters are located in the superior medial protocerebrum (mPME), lateral to the antennal lobe (mLALE), and in the gnathal ganglia (mVME) and project bilaterally to both AMMCs.

**Figure 1:**
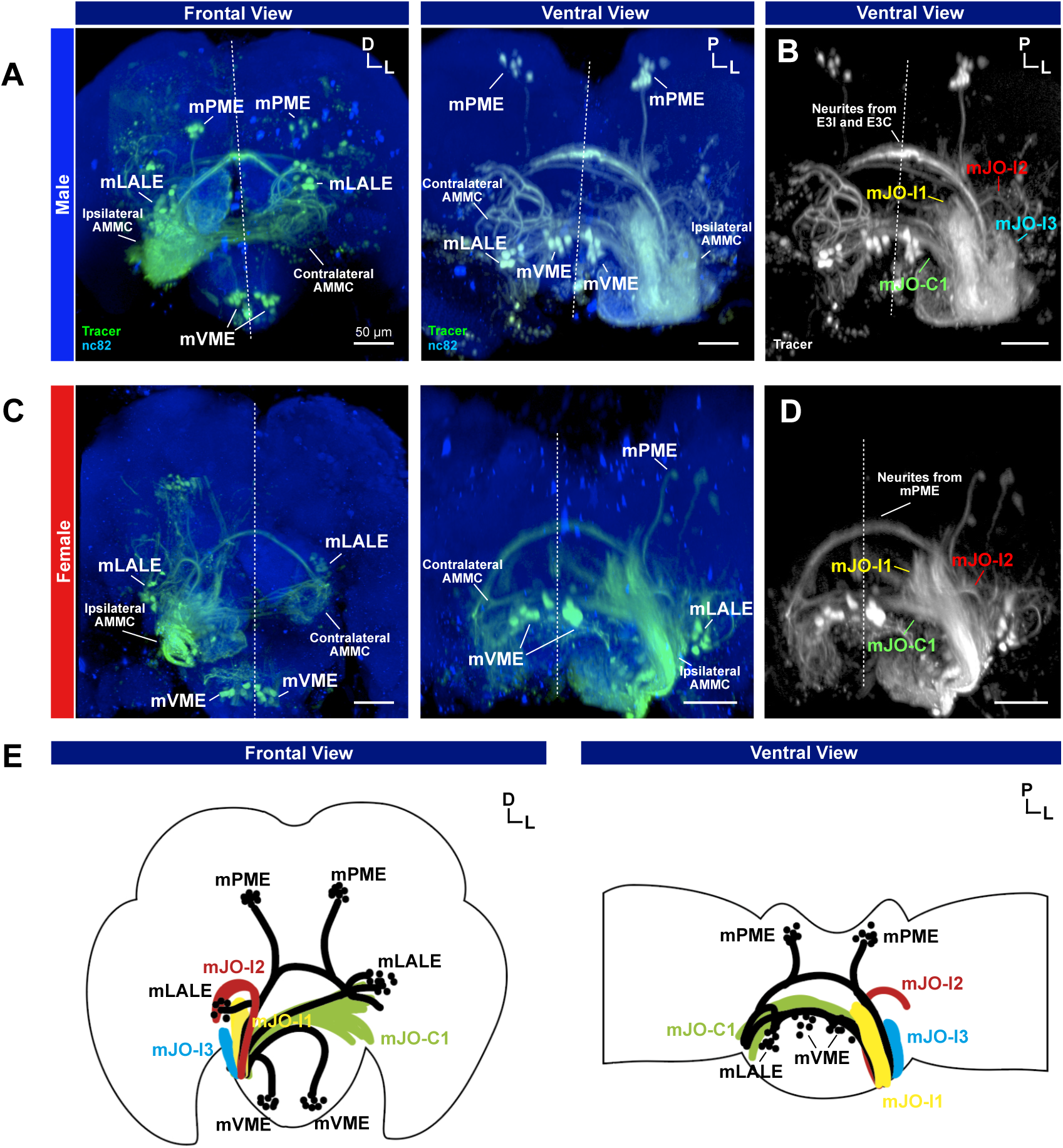
**Anatomy of JO neuron axons and efferent neurons in the brain.** (A to D) Confocal optical sections of mosquito brains with fluorescent tracer injections. JO neuron axons and efferent neurons in males (A, B) and females (C, D) are visualised (green or white). Scale bar = 50 µm. (A, C) Cell bodies of efferent neurons were labeled (mPME, mLALE and mVME). Ipsilateral and contralateral AMMCs correspond to the tracer-injected pedicel. Frontal (Left) and ventral (Right) views are shown. nc82 antibody labeling visualises neuropils (blue). Dashed line represents the brain midline. D, dorsal; L, lateral; P, posterior (the same in the following figures). (B, D) Axon bundles of JO neurons. mJO-I1-3 projected to the ipsilateral side while mJO-C1 innervated the contralateral side. (E) Schematics of labeled neurons in male brains. Left panel, frontal view; Right panel, ventral view. Coloured nerves represent JO neurons and black nerves show efferent neurons.

Axons projecting from the pedicel to the AMMC bifurcated into at least four branches in males and three branches in females (Fig. 1B, D). Axons in three branches projected to the ipsilateral brain (mJO-I1-3), and one bundle innervated the contralateral brain (mJO-C1). The medial branch, mJO-I1, directly extends to the posterior brain region. The shorter lateral branch, JO-I3, appeared only in males and resembled JO-A or JO-B axonal branch in *Drosophila*, which transmits sound information to higher-order neurons^37,42^. The intermediate branch, JO- I2, passed through the dorsal side of JO-I1 and innervated the dorsal posterior AMMC, drawing an arc.

Taken together, JO neurons in both sexes projected to both ipsilateral and contralateral AMMCs, although female JO neurons appeared to have fewer branches.

### Sexual dimorphisms of sound representation in the brain

*Ae. aegypti* male and female flagellae are mechanically tuned to different frequencies of sound^9^, with the male flagellum tuned to frequencies similar to female WBFs. Interestingly, male JO neurons have been reported as most sensitive to frequencies of sound significantly lower than female WBFs^9^. To reconfirm these findings and investigate the electrical tuning of female ears, we used a combined laser Doppler vibrometry/electrophysiology paradigm to simultaneously measure mechanical vibrations of the flagellum and antennal nerve compound action potential signaling in response to stimulation (Fig. 2A, B). We found mean mechanical tunings of 431 Hz and 253 Hz for male and female *Ae. aegypti* respectively, with mean electrical tuning estimates of 346 Hz and 220 Hz (Fig. 2B). Males thus have significantly higher mechanical and electrical tuning peak responses than females (p < 0.0001 for both mechanical and electrical), though both have higher mechanical than electrical tunings (p < 0.0001 for males and p = 0.0222 for females).

**Figure 2:**
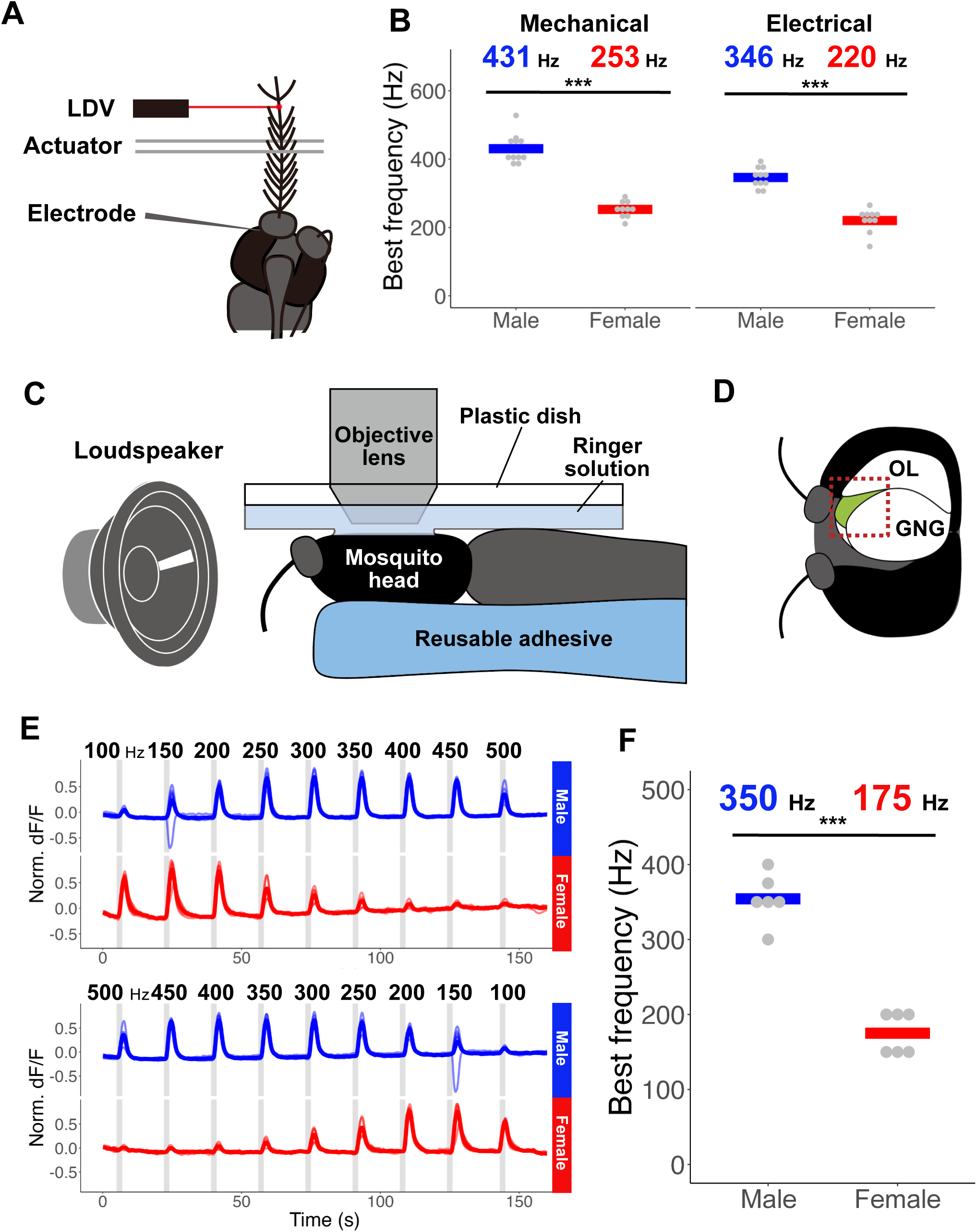
**Auditory response properties of JO neurons and AMMC.** (A) Experimental setup to measure mosquito ear mechanical tuning and electrical tuning. (B) Best frequencies of mechanical (Left) and electrical (Right) tuning. Dots and crossbars indicate the best frequency for each individual and mean of all individuals, respectively. *** p < 0.01 (ART ANOVA with Tukey method). (C) Experimental setup for calcium imaging. (D) Location of the AMMC in calcium imaging setup. The AMMC (Green area) can be observed between the optic lobe (OL) and gnathal ganglia (GNG). The red dotted square shows the region of interest used for the analysis. (E) Time traces of AMMC calcium responses to pure tones. Responses in males (top) and females (bottom) are shown. Pure-tone frequency was ramped up in top panels and decreased in bottom panels. Thin lines show time traces of the response for each individual. Bold lines indicate the average of all individuals. Norm. dF/F, normalized dF/F. (F) Best frequencies of the overall AMMC responses in males and females. Tone frequencies which evoked the maximum response are plotted. Dots and crossbars indicate the peak frequency for each individual and mean of all individuals, respectively. *** p < 0.01 (ART ANOVA).

WBF recordings of free-flying mosquitoes gave estimates of 499 Hz and 819 Hz for females and males, respectively (p = 2.2e-16; Supplementary Figure 2A). The estimated QDP in these conditions is thus 320 Hz, similar to the estimated peak electrical tuning of males of 346 Hz. We found peaks at lower frequencies as well which appeared similar in frequency to peak female electrical tuning responses, potentially indicating conservation of some neuronal populations between the sexes (Supplementary Fig. 2B). However, our methodology of recording electrophysiological responses from the entire antennal nerve bundle precluded the separation of potentially different populations.

Thus, we next used calcium imaging for further investigation of JO neuron responses to sound in greater detail. Given that JO neurons mainly project into the AMMC in both sexes, we investigated the frequency response properties of AMMC (Fig. 2C, D). We utilized the Q/QF system to drive fluorescent calcium reporter GCaMP6s expression pan-neuronally using a *brp-QF* driver line and observed sound-evoked calcium responses in the brain. We first delivered pure tones with a wide frequency range (50-850 Hz). In males, the AMMC strongly responded to 300-400 Hz of pure tones (Supplementary Fig. 2C). In contrast, the female AMMC showed strong responses to pure tones of around 100-200 Hz (Supplementary Fig. 2C). Building on this, we delivered individual pure tones focusing on a frequency range of 100 and 500 Hz to confirm the peak frequency, which was detected at 350 Hz in males and 175 Hz in females (p = 0.0001) (Fig. 2E, F). These estimates fit closely with peak frequency estimates from electrophysiology measurements (Fig. 2B).

### Mosquito AMMC has functionally distinct auditory regions

To distinguish functionally different neuronal populations, we analyzed the response properties of each pixel of our AMMC images (Fig. 3A). We first normalized time traces of fluorescent changes with maximum values to enable comparison of response patterns. To extract only reliably responsive pixels, we discarded pixels with weak or unstable responses (see Methods), yielding a 12,126 x 1,700 matrix (pixels x time). We then mapped the frequency to which each pixel showed the largest response to visualise peak frequency responses in the AMMC. Female AMMCs showed typical response peaks in the range of 100-200 Hz, while male responses were mainly restricted between 250-450 Hz (Fig. 3B). Unlike fruit flies ^42–45^, the spatial organization of peak frequency appeared divergent among individuals, with male AMMCs more diverse in peak frequencies than females.

**Figure 3:**
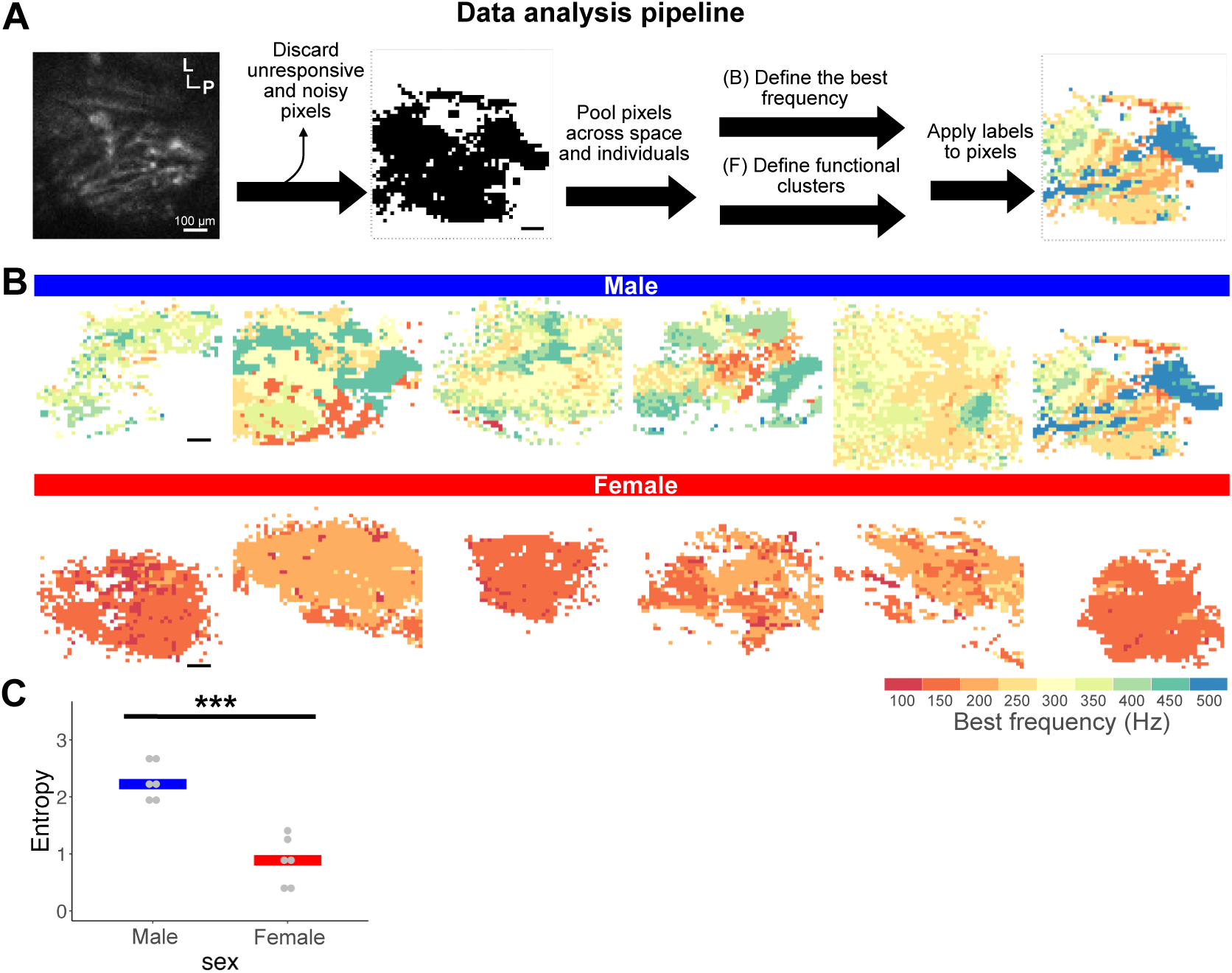
**Spatial variation in AMMC best frequency.** (A) Analysis pipeline. Fluorescent images of the AMMC are processed to obtain pixel images with colour-coded response properties. (B) Best frequency map of the AMMC. Maps from six males (top) and six females (bottom) are shown. Scale bar = 100 µm. (C) Diversity of best frequencies across space in males and females. The entropy across pixels in the AMMC is a measure of diversity. Dots and crossbars show the entropy for each individual and median of all individuals, respectively. *** p < 0.01 (ART ANOVA).

To validate sexual dimorphisms in peak frequency divergence, we calculated the entropy for each individual as a measure of frequency diversity. We obtained a significantly higher entropy value for males, indicative of a wider peak frequency range than for females (p = 0.0002; Fig. 3C) and agreeing with the multiple peaks covering a broad range of frequencies observed in the electrical responses of some male JOs (Supplementary Fig. 2B).

Next, we mapped representations of the whole response features by categorizing all waveforms among pixels (Fig. 4). Taking the ambiguous spatial organization into consideration, we split the response types into the smallest number of visually distinct clusters. When we set the threshold distance to 70, the response properties were categorized into 10 clusters (Fig. 4A, B). Male and female response properties were almost entirely distinguishable, with only one cluster (#7) present in both males and females. Six clusters (#1-6) were exclusively found in male AMMCs, and three clusters (#8-10) were observed only in females (Fig. 4A, C). Most male clusters tended to show strong responses to a wide range of frequencies. In contrast, female AMMCs responded to more specific frequencies (Fig. 4B). Male clusters include more diverse best frequencies, suggesting that a part of the diversity of best frequencies in male AMMCs was due to these wider response properties (p = 0.0255; Fig. 4D, E).

**Figure 4:**
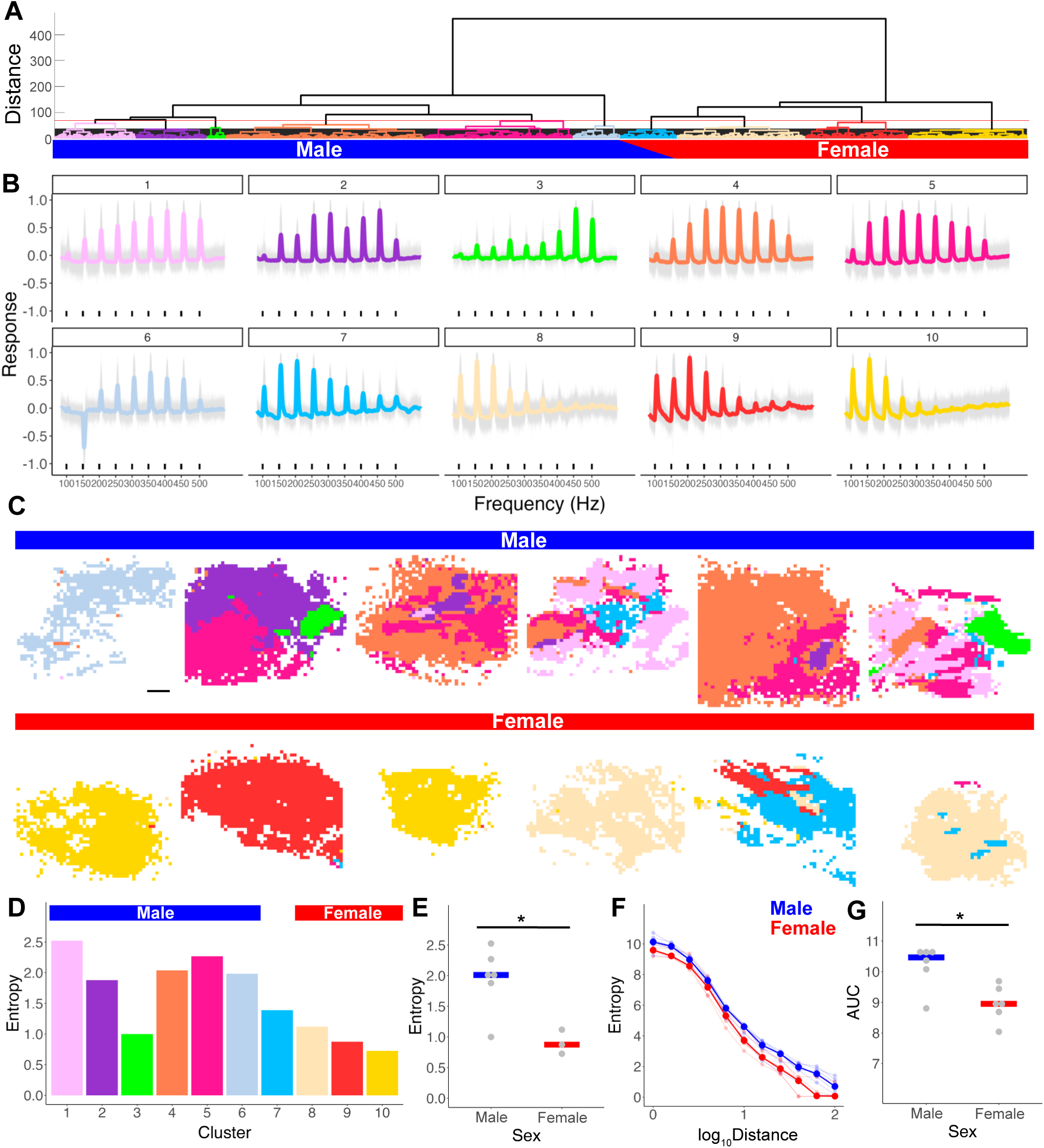
**Spatial variation of auditory representation in the AMMC.** (A) Dendrogram of hierarchical clustering. Pixels were divided into 10 functional clusters using a fixed threshold of 70 (red horizontal line). The same colour code for each cluster is used in panels B-D. (B) Response patterns of each cluster. Clusters 1 to 6 belong to males, cluster 7 to both males and females, and clusters 8 to 10 to females. (C) Spatial mapping of functional clusters in the AMMC for representative individuals. Scale bar = 100 µm. (D) The entropy of best frequencies across pixels (as a measure of how strongly each pixel responded to a wide range of frequencies) for each cluster. (E) Entropy of clusters appearing only in males and females. * p < 0.05 (ART ANOVA). (F) Entropy of functional types across space through various distance thresholds. (G) Area under the curve of (F). * p < 0.05 (ART ANOVA).

Although we tried to align the angle and depth of AMMC as precisely as possible, we could not find any common pattern among individuals (Fig. 4C). While female AMMCs show specific response patterns, male AMMCs appeared to have more functionally diverse pixels. Since the entropy of clusters across pixels was higher in males throughout the distance, we conclude that male AMMCs are more functionally diversified than females (p = 0.0158; Fig. 4F, G).

### Self-sustained oscillation may produce anti-resonance of a flagellum

Surprisingly, cluster #6 (identified only in males) showed a strong negative calcium response to 150 Hz (Fig. 4B, 5A), with at least two males displaying a drop in calcium concentration in response to stimulation of this frequency. One possible reason is that the flagellae of these males were vibrating in distinct states to other males, i.e., demonstrating SSOs^9^. We hypothesized that playback of a 150 Hz tone minimized the male flagellar oscillation whilst in the SSO state, causing the reduction of calcium level at 150 Hz.

To test this hypothesis, we investigated the frequency components of the male flagellar response to white noise (40-2000 Hz; Fig. 5B). First, we injected male mosquitoes with DMSO, which has been shown to induce SSOs^9^. Using laser Doppler vibrometry, we then monitored the response movement of the flagellum when exposed to the white noise.

**Figure 5:**
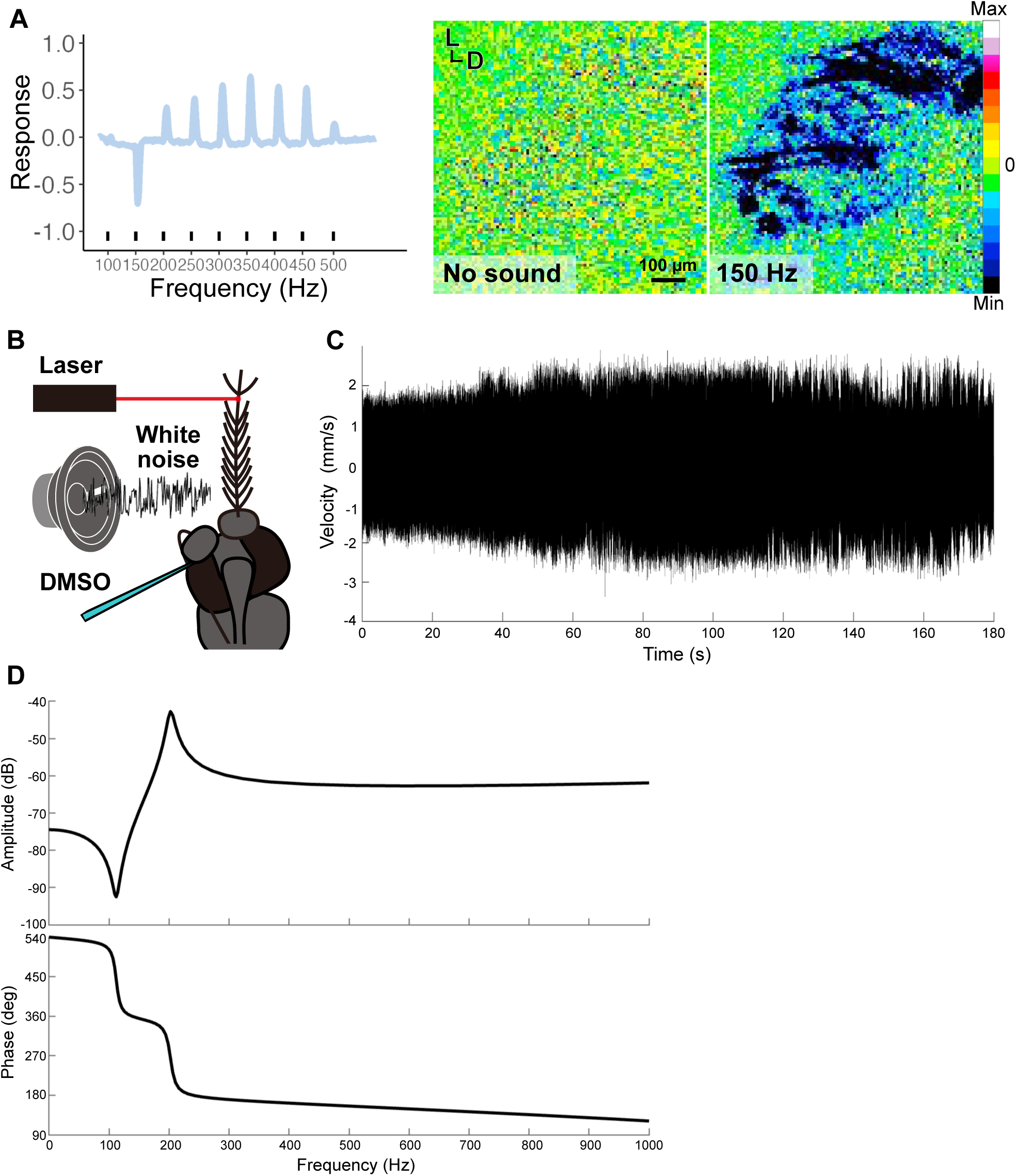
**Anti-resonance profiling in flagellum of males exhibiting SSOs** (A) Calcium response of cluster 6 shown in Figure 4B. Left panel, time trace of mean calcium response. Right panel, pseudocoloured changes in calcium level with no sound stimulus or during 150-Hz tone playback. (B) Experimental setup of laser Doppler vibrometry. To induce SSOs in males, DMSO is injected to the base of the other pedicel. (C) Example of velocity time trace of flagellar vibration during white noise exposure. (D) State-space modeling of (C). Amplitude (Top) and phase (Bottom) of flagellar vibration across frequencies.

The continuous changes in the envelope of time traces of flagellar vibrations encouraged us to use quadratic state-space modeling for our simulations (Fig. 5C). Three of twelve male SSO flagellae demonstrated a negative peak with an anti-resonant point around 150 Hz (Fig. 5D, Supplementary Fig. 3A). In the quiescent state (i.e. non-SSO state), on the other hand, no anti-resonance was found in any individuals (Supplementary Fig. 3B). This suggests that ∼150 Hz tones can minimize flagellar vibrations in some cases, bringing about a drop in calcium level in specific individuals (cluster 6).

### Cilium-related genes are highly enriched in male pedicels

To investigate the molecular basis of sexual dimorphisms in auditory responses in the AMMC, we compared between sexes the gene expression profile of the pedicel, which houses JO including cell bodies of JO neurons and support cells^46^. We first verified the quality of RNA sequencing with principal component analysis (PCA) and found that most of the observed variance was attributable to tissue (43%) and sex (20.7%), indicating that biological replication effects were relatively negligible (Fig. 6A). DESeq2 analysis identified 820 transcripts enriched in male pedicel compared to female pedicel, and conversely, 1377 transcripts were found to be more abundant in female compared to male pedicels (p < 0.1, log_2_ fold change > 1, Fig. 6B, C).

**Figure 6:**
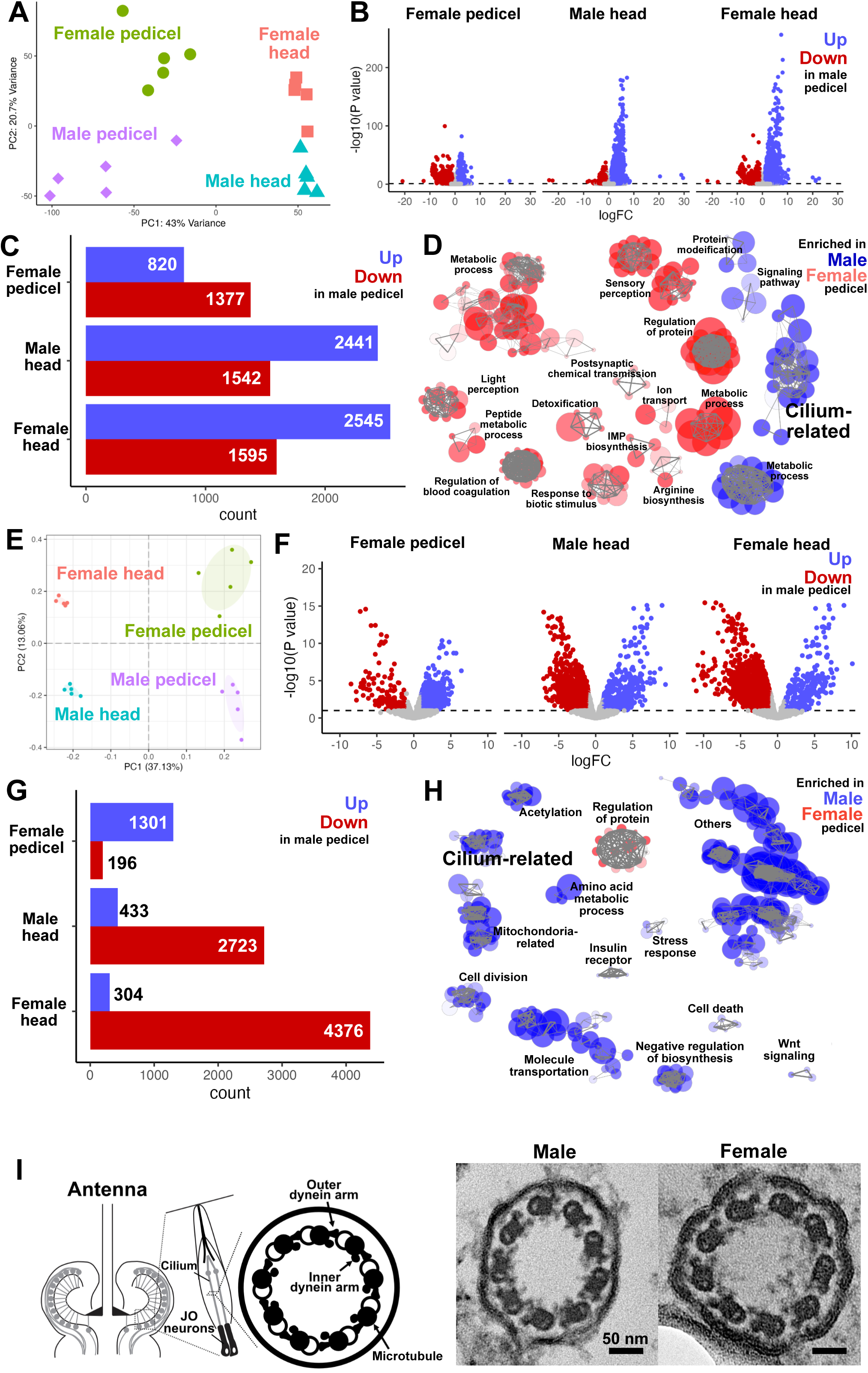
**Transcriptome and proteome analysis.** (A to D) Transcriptomic analysis of pedicels and heads in males and females; five replicates per group. (A) Principal component analysis. (B) Volcano plot of expression differences between male pedicels and others. Genes with adjusted p value < 0.1 and log fold change (logFC) > 1 were coloured. (C) Differentially expressed genes between male pedicels and others. (D) Enrichment GO network analysis. GO terms of genes enriched in males (blue) and females (red) were displayed as nodes. Colour intensity and size of nodes represent adjusted p value and number of genes respectively. Thickness of lines indicates the percentage of overlapping genes. GO terms with adjusted p value < 0.1 and more than two edges were described. Each GO term can be found in Supplementary file 1, 2. (E to H) Proteome of pedicels and heads in males and females; five replicates per group. (E) Principal component analysis. (F) Volcano plot of expression difference between male pedicels and others. Genes with adjusted p value < 0.1 and log fold change (logFC) > 1 were coloured. (G) Differentially expressed genes between male pedicels and others. (H) Enrichment GO network analysis. Individual GO terms can be found in Supplementary files 3 and 4. (I) Structure of cilium in JO neurons. Left panel, the diagram of cilium structure in JO neurons and a pedicel. Right panel, Transmission electron microscopy images of cilium in male and female JO neurons. Scale bar = 50 nm.

Gene Ontology (GO) enrichment analysis identified GO terms annotating upregulated and downregulated genes in male pedicels than females (Table1, Supplementary table 1). In these GO terms, gene overlaps were frequently observed in both sexes (Tables 1, 2). To visualise GO terms that label the same genes, we measured the distance among terms by the proportion of shared genes and constructed the network (Fig. 6D, Supplementary Fig. 4). All GO terms of upregulated genes in the male pedicel were clustered into four clusters, one of which was related to the construction of cilium. Female pedicel-enriched genes comprised of 13 clusters, including regulation of blood coagulation and biotic stimulus detection.

To test if the upregulation of cilium-related genes was also detected at a protein level, we next conducted LC/MS-MS proteomic analyses of mosquito ears and heads. PCA analysis showed that over 50% of the variance could be ascribed to tissue (37.13%) and sex (13.06%), validating that the replication effects were negligible also in proteome analysis (Fig. 6E). Linear model analysis identified 1301 proteins enriched in male pedicel compared to females (p < 0.1, log_2_ fold change > 1, Fig. 6F, G). GO network analysis confirmed that cilium-related proteins were also upregulated in male pedicel in comparison to females (Fig. 6H). Together, we concluded that molecules related to cilium construction are more highly expressed in male pedicels than female pedicels.

The cilium is located at the distal dendrite of JO neurons, encouraging some researchers to hypothesize that the cilium is a source of energy for the self-sustained flagellar oscillations (SSOs) found only in males (Fig. 6I)^19,26,27^. We next examined the effect of sexual dimorphisms in the expression of cilium-related genes on the cilium structure of JO neurons. Analysis using transmission electron microscopy (TEM) confirmed that both sexes shared a 9x2+0 structure of axonemal cilium as previously described^26,28^. We also found inner and outer dynein arms associated with microtubule doublets in both sexes, suggesting that both sexes share the basic structure of axonemal cilium in their JO neurons despite differences in cilium-related RNA and protein expression (Fig. 6I).

## Discussion

Mosquitoes exhibit a variety of sex-specific behaviours, making them a suitable model for investigating mechanisms of sexual dimorphisms in behaviour^39,47^. Here, we compared the auditory response properties of the brain between males and females and identified sex-specific spatial diversity in the auditory representation of the primary auditory center, AMMC. We also investigated molecular differences in the hearing systems of males and females using a combination of gene-expression profiles of hearing organs and electron microscopy.

Fluorescent tracer injection into the pedicel allowed us to trace whole neurons connecting to the pedicel, labeling both JO neurons and putative efferent neurons. Their neurites overlap with each other in the AMMC, precluding us from describing the detailed bifurcation pattern of JO neurons. However, the fluorescent tracer labeled at least four axon bundles of male JO neurons and three branches in females. A branch of JO neurons innervates the contralateral AMMC, in contrast to fruit flies whose JO neurons do not project to the contralateral AMMC^37,41^. As for optic-nerve projection through the optic chiasm in vertebrate visual pathways, signals detected by one ear might therefore be partially transmitted to the contralateral AMMC in mosquitoes^48^.

One possible function of this binaural convergence is to represent the signals from both ears for encoding acoustic field representation, equivalent to visual field representation in the vertebrate brain to enable stereopsis. It should be noted, however, that the JO not only detects sound but also gravity and wind in *Drosophila*^42,44^. Further investigation is needed to elucidate if and how the mosquito AMMC is divided into functional subregions, and how the information delivered from both ears is interactively processed in the AMMC.

In this study, the tracers injected into the gap created by removing both the flagellum and the hole made by removing the pedicel labeled the AL. A previous study reported that mosquito JO neurons project to the subset of AL known as the JO-C (JO-Centre) where auditory and olfactory information might be integrated^38^. Our neuro-tracing experiments lend support to this previous report, although the tracer from the pedicel may also have labeled olfactory sensory neurons originating from the flagellum. Calcium imaging of the AL would help test whether auditory information is represented also in the AL.

Previous studies in humans have reported sexual dimorphisms in the cochlea, an equivalent hearing organ to the mosquito JO, in terms of size and stiffness gradient^49–52^. However, large sex differences in frequency properties are rarely observed in the early stages of auditory processing across a wide array of animals, including humans (but see Krizman et al^50^)^53–55^. In the present study, we found that the frequency characteristics of the AMMC, the first stage of auditory processing in the mosquito brain, are significantly distinct between the sexes, suggesting that strong sexual selection pressure on mosquitoes has brought about these substantial dimorphisms.

Male AMMCs showed peak responses to 350 Hz of sound, similar to the estimated peak electrical tuning of JO neurons estimated via electrophysiology (∼346 Hz) and similar to the best frequency of the QDP (320 Hz). Our study thus lends support to the DP-based communication hypothesis in mosquito acoustic communication, and opens new avenues for further understanding the neural basis of their acoustic communication. The peak responses of the female AMMCs were lower than the estimated peak electrical tuning of their JO neurons (∼150 Hz compared to 220 Hz).

Considering the distinct behaviours observed in male and female mosquitoes, sexual dimorphisms in the information processing pathways of various sensory modalities has likely evolved to support their sex-specific behaviours, such as phonotaxis, blood-feeding and host-seeking. This is in part reflected in changes in brain morphology, with the AMMC significantly larger in males than females, and the AL significantly larger in females than males^39^. The analytical tools we developed in this study should help identify the sex-biased landscape of sensory processing in the mosquito brain.

Our state-space modeling suggested that SSOs can interfere with the flagellum response at 150 Hz tone, bringing about the negative calcium response of the AMMC in males. We measured the vibration of SSO flagellar induced by DMSO injection, which causes non-specific physiological impairments. The anti-resonance of flagellar appeared only in three of twelve individuals. It raises two questions. Firstly, why was anti-resonance not found in all DMSO injected mosquitoes? One possible reason is that DMSO damaged not only the suppression system of SSOs but also the key mechanism of active hearing in the other mosquitoes. Secondly, what is the biological role of anti-resonance in SSOs? SSOs appeared so rarely that no physiologically specific induction of SSOs was reported in *Ae. aegypti*^9,22^. SSOs entrain only to pure tones similar to female WBFs, suggesting that the anti-resonance produced by SSOs is a byproduct of maximizing sensitivity to female flight tones. Calcium imaging of auditory responses of SSO-induced brains would support our understanding of how SSOs alters auditory processing in the brain^48^.

Our transcriptome and proteome analyses revealed enrichment of GO terms related to the regulation of blood coagulation and responses to biotic stimulus in females, potentially relevant for blood aspiration. On the other hand, microtubules, dynein and related genes (i.e., cilium-related genes) were found to be highly enriched in the male pedicel. However, electron- microscopy images suggest that both males and females share the basic cilium structure of the JO neuron’s dendrite. Males and females may therefore potentially express different sets of cilium-related genes with distinct kinetic properties, enabling specific flagellar oscillation. Calcium imaging of males whose male-specific cilium-related genes are knocked down may help our understanding of the role of the male-specific gene set in the male-specific response properties in the AMMC.

## Methods

### Mosquito rearing

All *Ae. aegypti* mosquitoes were reared using a 12 h:12 h light–dark cycle at 28 °C and 60-70% relative humidity. Adults were fed a 10% glucose solution^56^. For calcium imaging, we used *brp-QF2w*^56^ as a driver line and *QUAS-GCaMP6s*^56^ as a reporter line. The wild-type *Liverpool* strain was used for all other experiments.

### Tracer injection

Mosquitoes three to ten days after eclosion were anesthetized on ice and one of their flagellae was clipped from the pedicel. To visualise the nerve from the flagellum, a drop of fluorescent tracer (Dextran, Tetramethylrhodamine, and biotin, 3000 MW, Lysine Fixable; D7162, Invitrogen) dissolved in phosphate-buffered saline (PBS; #T9181, Takara) was placed onto the gap created by removing flagellum. Mosquitoes were then kept at 4 °C for an hour. To visualise the nerve from the pedicel, the pedicel whose flagellum was clipped was then removed on ice and a drop of Alexa Fluor™ 488 (Dextran, 10,000 MW, Anionic; Thermo Fisher Scientific), which does not cross gap junctions, was placed onto the hole. The mosquitoes were then kept at 4 °C for three hours.

Mosquito heads were fixed in 4% paraformaldehyde (PFA) in phosphate-buffered saline (PBS) at 4 °C for three hours. The brains were dissected from the head in PBS, kept at 4°C in PBS containing 0.5% Triton X-100 (PBT) overnight, and incubated with primary (Mouse anti-Bruchpilot nc82) and secondary (Alexa Fluor 647-conjugated anti-mouse IgG) antibodies for 3 days respectively at 4 °C. After rinses with PBT and PBS, brains were incubated in 50% glycerol in PBS for an hour and 80% glycerol in deionized water for 30 min. Brains were mounted on glass slides for imaging.

### Confocal microscopy and image processing

Confocal images of brains were obtained at 0.84-µm intervals with an FV 1200 laser-scanning confocal microscope (Olympus, Tokyo, Japan) equipped with a silicone-oil immersion 30x Plan-Apochromat objective lens (NA = 1.05).

### Electrophysiology: recordings

Electrophysiological recordings utilised two-days entrained mosquitoes at Zeitgeber time (ZT) 11- ZT 13, in a temperature-controlled room at 25 ± 1.5 °C. Mosquitoes were mounted on rods using glue such that only their right flagellae were free to move. The rod was then placed in a micromanipulator on a vibration isolation table.

A reference electrode was inserted into the mosquito thorax to facilitate charging to -30 V relative to ground. Electrostatic actuators were placed around the freely moving flagellum so that electrostatic stimulation could be provided. The insertion of a recording electrode into the base of the pedicel enabled electrical recordings from the antennal nerve. A laser Doppler vibrometer was focused on the tip of the flagellum to record flagellar displacements to stimuli.

Next, a calibrated force-step stimulus was provided to calibrate the displacement of the flagellum to approximately ±3.5 μm; the stimulus was provided using a data acquisition unit CED Micro1401 (Cambridge Electronic Design). Following this calibration, sweep stimuli consisting of 1-second-long chirps of linearly increasing or decreasing frequency (i.e., 1 to 1000 Hz or 1000 to 1 Hz), were provided. Each loop consisted of 10 sets of sweeps, each containing 4 different sweeps: forward phasic, forward anti-phasic, backward phasic and backward anti-phasic. Between phasic and anti-phasic sweeps 0.1 s of silence was provided, whilst 0.4s of silence was provided between forward and backward sweeps. Flagellum displacement and nerve recording data were recorded using the Spike2 software (Cambridge Electronic Design).

### Electrophysiology: data analysis

For mechanical tuning analyses to calculate the flagellar tuning frequency, only data for which the laser quality data was recorded as greater than zero was included. A DC remove with time constant of 0.01 s was first applied to the laser data, before rectification and a smooth function with time constant of 0.0005 s. A slope function with 0.1 s time constant was used to find when the channel gradient was equal to zero; this represents the timepoint of maximum flagellar vibration. Calculation of this time allowed for estimation of the corresponding stimulus frequency, hereafter referred to as flagellar ear mechanical tuning frequency.

For electrical tuning analyses, nerve responses from sequential phasic and anti-phasic stimuli were averaged to cancel artefacts recorded in the nerve channel resulting from electrostatic actuation. This processed nerve signal was analysed as described above for the laser signal. DC remove, rectification and smooth functions were applied, followed by a slope function. Estimating the time at which the slope equaled zero enabled calculation of the corresponding stimulus frequency, denoted as the ear electrical tuning frequency.

In total, 11 male and 10 female electrophysiology recordings were collected and analysed.

### Mosquito WBF measurements

Groups of 50 virgin male or female Liverpool mosquitoes were entrained in incubators for two days at 25 ± 1.5 °C. During dusk on the third day, a microphone (EK series, Knowles) was inserted into the cage and flight tones recorded during the period immediately preceding dusk using a Picoscope 2408B and the Picoscope 6.14 software (Pico Technology) at a 50 kHz sampling rate. Data was analysed using the SimbaR package in R with only flight recordings longer than 150ms included in the final analysis.

In total, WBF recordings were collected and analysed from 11 male and 12 female cages.

### Calcium imaging: data acquisition

Mosquitoes were collected within 24 h after eclosion and less than eight mosquitoes of each sex were separately kept in individual vials. They were maintained for five to nine days after eclosion. Mosquitoes at the ZT 10-13 were anesthetized on ice and stabilized ventral side up onto an imaging plate using UV-cured adhesive (Norland Optical Adhesive 81, Norland). The head angles were adjusted to prevent flagellum from dipping into the Ringer’s solution and the dorsal head and thorax were glued with the adhesive to fix the flagellum direction. The posterior head and thorax were also glued to avoid destroying the cap structure during the following step. The ocelli of medial head and the mouthpart were removed using an incision scalpel (MICRO FEATHER®, Feather) and forceps to open a window to monitor GCaMP fluorescence from the AMMC in the brain. A drop of an adult saline solution with the following components was added to prevent dehydration: 108 mM NaCl, 5 mM KCl, 2 mM CaCl_2_, 8.2 mM MgCl_2_, 4 mM NaHCO_3_, 1 mM NaH_2_PO_4_, 5 mM trehalose, 10 mM sucrose, and 5 mM HEPES, pH 7.5, 265 mOsm. A fluorescent microscope (Axio Imager.A2, Carl Zeiss) equipped with a water-immersion 20× objective lens (W Achroplan/W N-Achroplan, NA = 0.5; Carl Zeiss), a spinning disk confocal head CSU-W1 (Yokogawa Electric Corporation), and an OBIS 488 LS laser (Coherent) for excitation at 488 nm was utilized.

To provide sound stimuli, a loudspeaker (FF225WK, Fostex) was positioned ∼11 cm from the antenna of the mosquito. For monitoring calcium responses to pure tones, a series of pure tones with different frequencies (50-850 Hz by 50 Hz for Supplementary Figure 2 and 100-500 Hz by 50 Hz for Figure 2-4) were delivered. The peak-to-peak amplitude of the tone was ∼5 mm/s for each pure tone.

### Calcium imaging: analysis

Calcium imaging data was analyzed using MATLAB (MathWorks), Fiji, and R software. Motion artifacts due to animal movement were cancelled using the NormCorre algorithm by running the CaImAn toolbox with MATLAB. For Supplementary Figure 2, the region of interest (ROI) was set at the bulging area in the lateral anterior brain. The mean fluorescence intensity in the ROI for each frame was calculated using Fiji. Moving averages of fluorescence intensity for three frames were adopted for data analysis to acquire smooth-line traces of calcium fluorescence. For Figures 2-4, the 50 x 50 pixels around the AMMC were selected, and responsive pixels were extracted using the following process. The relative fluorescence change (*dF/F*) was calculated to indicate the calcium response, which was given by

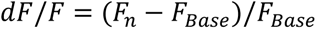

where *F_n_* is the corrected fluorescence intensity at n seconds from the sound onset. *F_Base_* is the average of the corrected fluorescence intensity during the 5 s before the stimulus onset. The noise (>3 kHz) of *dF/F* was cut with a Butterworth filter. The filtered value was normalised by the maximum value of each trial (normalized *dF/F*), allowing us to compare the response properties with the following process. The non-responsive pixels and highly dispersed pixels among trials were discarded to ensure the quality of analysis. Time trace of calcium response in each individual was averaged across pixels for Figure 2, and fluorescent movement in each pixel was analysed in Figures 3-4. The best frequency was identified based on the timing of the maximum fluorescent value. The hierarchical clustering was conducted based on Euclid distance, and the cutoff criterion was set at 70.

### Laser Doppler vibrometry with DMSO injection

Male mosquitoes were prepared as for the aforementioned electrophysiology experiments at 25 ± 1.5 °C. The laser Doppler vibrometer was focused on the mosquito’s right flagellum as described above, and 50% DMSO in Ringer’s solution was injected into the base of the mosquito’s left pedicel. White noise playback (1-2000 Hz at a Particle Velocity Level of 1.58 x 10^-4^ ms^-1^, equivalent to Sound Pressure Level of 70 dB) was provided using a loudspeaker (FF85WK, Fostex). Flagellar velocity in response to white noise stimulation was measured for the next 3 min at a sample rate of 12 kHz using the VibSoft 6.0 software (Polytec).

### Modeling of antenna vibration by State-Space method

The behaviour of an antenna when it receives white noise stimuli was modeled using the state-space method in control theory. The number of state variables needs to be determined when expressed in state space, and, by observing the behaviour of the antenna in Figure 5C, it can be assumed to be a vibration phenomenon with multiple frequency components. In general, this vibration phenomenon can be modeled as a coupled oscillation caused by connecting multiple springs. When we model antenna vibration, one side is assumed a fixed end, and the other side is a free end, because the antenna is fixed to the head and the other tip is free. The simplest coupled oscillation can be formulated as follows:

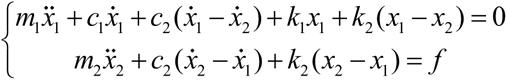

where, m, c, and k represent the coefficient of the mass, damper, and spring that make up the coupled oscillation system, respectively, and the subscripts 1 and 2 denote the number of components. The minimum coupled oscillation system has two degrees of freedom and consists of two masses, springs, and dampers. This time, the equation of motion is derived with subscript 1 as the fixed end. The following state equation can be obtained by modeling this simultaneous differential equation as a state space in control theory.

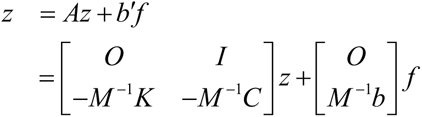

where the elements of each matrix are defined as follows:

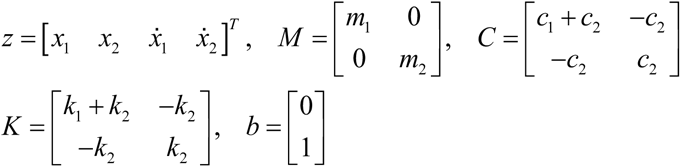

In other words, when modeling the behaviour of an antenna using the state space method, we can model it in a state space with four state variables. The state space with four state variables was identified based on the relationship between the white noise sound input waveform and the behaviour of the antenna as output obtained in the experiment.

The identification of the variables was performed using the System Identification toolbox of MATLAB.

### Transcriptomics: sample preparation

Groups of virgin Liverpool male or non-blood fed female mosquitoes were flash-frozen in liquid nitrogen at ZT12 after entrainment in an incubator for two days. Tissues were dissected on ice in RNAiso Plus (#9109, Takara Bio Inc.). Five repeats of pedicels and heads were collected, and RNA extracted from homogenized samples. RNAiso Plus was added to lysed samples for a final volume of 1 ml. Samples were incubated at room temperature for 5 min before 200 µl chloroform (Kanto Chemical Co., Inc.) was added prior to mixing. Following a further 15 min incubation at room temperature, samples were centrifuged at 12,000 × g for 15 min at 4°C. 0.5 ml 2-propanol (Sigma) was added, and samples were stored for 30 min at −20°C.

Supernatant was discarded following centrifugation at 12,000 × g for 10 min at 4°C, and the remaining RNA pellet washed with 1 ml 75% ethanol (Sigma) solution. Samples were centrifuged for 5 min at 7,500 × g at 4°C, with this ethanol wash cycle performed twice before a further wash with pure ethanol (Sigma). The RNA pellet was next dissolved in Nuclease Free Water (Invitrogen) and RNA quality assessed using a Nanodrop (ThermoFisher Scientific). RNA was then submitted to the Center for Gene Research at Nagoya University.

### Transcriptomics: analysis

The sequenced reads were aligned by using Bowtie2 and more than 90% of reads were aligned. The number of alignments mapped to each gene was counted with a gene transfer format (GTF) file of the AaegL5.0 genome from vectorbase (https://vectorbase.org/vectorbase/app). The read count data was analysed via “iDEP 2.0”^57^. Differentially expressed genes (DEGs) were analysed using DEseq2 and the GO terms for GO enrichment analysis were obtained from Ensembl Metazoa (LVP AGWG (aalvpagwg_eg_gene) dataset). The clusters in Figure 6 D, H were labeled by summarizing GO terms in the clusters and each GO terms can be confirmed in Supplementary File 1-12.

### Proteomics: sample preparation

Groups of virgin male and female mosquitoes were collected as for the RNA sequencing experiments described above.

Tissues were then dissected in PBS and transferred to microfuge tubes containing PBS at room temperature. PBS was replaced by sample buffer (100 mM Tris-HCl pH 8.0, 1% SDS, 20 mM NaCl) and tissues were homogenized on ice before being centrifuged at 16,100 g for 10 min at 4°C to remove debris. The extracted supernatants were transferred to new microfuge tubes, with 2 µl of supernatant per sample used to test protein concentration (Pierce 660 nm Protein Assay, Invitrogen, Cat: 22662). Samples were stored in a -20°C freezer prior to LC/MS-MS. Extracts were reduced with 25 mM dithiothreitol for 30 min at 25 °C, followed by alkylation with 50 mM iodoacetamide for 30 min at 25 °C in darkness.

Protein purification and digestion were performed using the sample preparation (SP3) method^58^. The beads were then resuspended in 100 μL of 50 mM Tris-HCl (pH 8.0) with 1 μg of Lys-C (Fujifilm-Wako) at 37 °C for 4 h followed by digestion with 0.2 μg of trypsin (Promega) and mixed gently at 37 °C overnight. The digested samples were acidified with 5 μL of 20% trifluoroacetic acid (TFA). Digested peptides were desalted using GL-Tip SDB (GL Sciences, Tokyo, Japan), evaporated in a SpeedVac concentrator, and redissolved in 0.1% TFA and 2% acetonitrile.

### Data-independent LC-MS/MS analysis

LC-MS/MS analysis of the resultant peptides was performed on an UHPLC connected to a Q-Exactive Orbitrap mass spectrometer (Thermo Fisher Scientific) through a nano electrospray ion source (AMR Inc., Tokyo, Japan). The peptides were separated on a 125-mm C18 reversed-phase column with an inner diameter of 100 µm (Nikkyo Technos, Tokyo, Japan) with a linear 5–40% acetonitrile gradient for 0–100 min, followed by an increase to 95% acetonitrile for 5 min. The mass spectrometer was operated in data-independent acquisition mode. The MS1 spectra were collected in the range of m/z 500–740 (isolation window width, 10 Da) at 70,000 resolution to set an automatic grain control (AGC) target of 3 × 106. MS2 spectra were collected at 200–1800 m/z at 70,000 resolution to set an AGC of 3 × 106, a maximum injection time of “auto”, and stepped normalized collision energies of 22%.

### Protein Identification and Quantitative Analysis from MS Data

DIA-MS files were searched against an *in silico* human spectral library using DIA-NN (version: 1.8.1). First, a spectral library was generated from the VectorBase-62_AaegyptiLVP_AGWG_AnnotatedProteins database using DIA-NN^59^. Parameters for generating the spectral library were as follows: digestion enzyme, trypsin; missed cleavages, 2; peptide length range, 7 to 45; precursor charge range, 1 to 4; precursor m/z range, 495 to 745; and fragment ion m/z range, 200 to 1800. “FASTA digest for library-free search/library generation” “deep learning-based spectra, RTs, and IMs prediction” “N-term M excision” and “C carbamidomethylation” were enabled. DIA-NN search parameters were as follows: mass accuracy, 10 ppm; MS1 accuracy, 10 ppm; protein inference, genes; neural network classifies, single-pass mode; quantification strategy, robust LC (high precision); and Cross-run normalization, RT-dependent. “Use isotopologues,” “heuristic protein inference,” and “no shared spectra” were enabled. Matching between runs was turned on for quantitative analysis. Protein identification threshold was set at 1% or less for both precursor and protein false discovery rates (FDRs).

### Proteomics: analysis

Most steps of analysis were conducted with “Bioconductor” packages^60^ in R software. The DEG was analysed with “limma-voom”^61^ and GO enrichment analysis was conducted using “iDEP 2.0”^57^.

### Electron microscopy

Mosquitoes were decapitated, and the proboscis and flagellum were clipped from the head in phosphate-buffered saline (PBS). Heads were fixed overnight at 4 °C in a fixative containing 2% EM grade paraformaldehyde, 2.5% glutaraldehyde and 0.08 M cacodylate. Heads were washed in the fixative three times and post-fixed in 2% OsO4 for three hours. After washing in the fixative, heads were dehydrated by immersion in increasing concentrations of ethanol solutions, up to and including pure anhydrous ethanol. To infiltrate with resin, heads were passed through pure propylene oxide (PO) for 10 min, 1:1 PO/resin for 4 h and pure resin for a day. Samples were then cured in silicon molds at 60 °C for 48 hours. Pedicels from fixed heads were sectioned using an ultramicrotome (UC7k, Leica) with diamond knives (ultra 35° and histo, DiATOME). Thin sections were collected, stained with 2% uranyl acetate and lead stain solution (Sigma Aldrich), and imaged using a Transmission Electron Microscope (JEM- 1400PLUS, JEOL).

## Acknowledgments

We thank Mika Nomoto, Yasuomi Tada, Akiko Akama, and Mikako Yamaguchi (Center for Gene Research, Nagoya University) for assistance with RNAseq experiments at the Division for Medical Research Engineering. We appreciate Koji Itakura and Nagoya University Graduate School of Medicine for assistance and use of the Ultramicrotome and TEM equipment. We also acknowledge Conor McMeniman for providing mosquito strains tested in this paper.

## Contributions

T.S.O., M.P.S. and A.K. conceived the study. T.S.O. and Y.N. performed neurotracing experiments. Y.M.L. and Y.Y.J.X. performed electrophysiology experiments. D.F.E. and M.P.S. analysed electrophysiology data. T.S.O. performed calcium imaging experiments. Y.Y.J.X. measured SSO vibrometry. T.S. and S.S. analysed calcium imaging data and SSO vibrometry data. T.S.O., Y.M.L. and M.P.S. performed transcriptomics experiments. T.S.O analysed transcriptomics data. E.M.S., T.S.O., T.L., and M.P.S. performed proteomics experiments. T.S.O. and E.M.S. analysed the proteomics data. T.S.O. and D.F.E. performed transmission electron microscopy experiments. M.P.S. and A.K. supervised the project. All authors contributed to paper writing.

## Data availability

All data is available via the following Dryad link

---

**Table 1: Gene ontology enrichment analysis.**

Gene Ontology (GO) terms (biological processes) and annotated upregulated and downregulated genes (adjusted p value < 0.1) in male pedicels compared to females.

**Table 2: Genes with multiple GO term annotations.**

Top 10 upregulated and downregulated genes in male pedicels compared to females based on the highest number of shared GO terms.

## Supplementary figure legends

**Supplementary figure 1: Axons of neurons projecting from pedicel and flagellum** Confocal optical sections of mosquito brains with fluorescent tracer injections. Neurons connecting to pedicel (green) and flagellum (magenta) in males (A) and females (B) are visualised. White dashed line represents the brain midline. D, dorsal; L, lateral; Areas surrounded by an orange dashed rectangle in left panels are magnified in the right panels.

**Supplementary figure 2: Wing beat frequencies, mechanical and electrical tuning, and calcium responses to a wide frequency range**

(A) Wing beat frequencies (WBFs) of male and female mosquitoes at 25°C. Dots and crossbars indicate the median WBF for each individual cage tested and mean of all individuals, respectively. *** p < 0.001 (Welch’s t-test). (B) Mechanical (Top) and electrical (Bottom) responses for males and females in response to sweep stimulation. Left panel, female; Middle panel, male with identifiable low-frequency peaks; Right panel, male without identifiable low-frequency peaks. Arrowhead indicates low-frequency peak. (C) Time traces of calcium responses during playback of tones covering wide range of frequencies. Top panel, responses to sweep-up tones; Bottom panel, Sweep-down tones. Thin lines represent individual responses and bold lines indicate mean values across individuals. Gray shaded areas indicate the times when sound stimuli were delivered.

**Supplementary figure 3: State-Space modeling of flagellar vibrations during white noise exposure**

State-space modeling of flagellar vibrations during white noise playback during (A) SSO and (B) normal states. Top panel, amplitude of vibration. Bottom panel, phase of vibration. (A) Four types of vibrations appeared. Anti-resonance around 150 Hz (Type 1) was found in three of the twelve individuals tested. (B) Normal state shows two types of flagellar vibrations. No anti-resonance was found in all individuals tested.

**Supplementary figure 4: Enrichment GO network of male pedicel vs male or female heads**

Enrichment GO network analysis for transcriptomic (A, B) and proteomic (C, D) analysis. (A, C) GO terms of genes enriched in male pedicels (blue) and male heads (red) were displayed as nodes. (B, D) GO terms enriched in male pedicels (blue) and female heads (red) were shown as nodes. Colour intensity and size of nodes represent adjusted p value and number of genes, respectively. Thickness of lines indicates the percentage of overlapping genes. GO terms with adjusted p value < 0.1 and more than two edges were described. Enrichment of cilium-related genes in the male pedicel are detected in all cases. Individual GO terms can be found in Supplementary files 5-12.

**Supplementary file 1: Interactive network of enrichment GO analysis of upregulated genes in male pedicel compared to female pedicels based on transcriptomic analysis (Related to Figure 6D).**

**Supplementary file 2: Interactive network of enrichment GO analysis of upregulated genes in female pedicels compared to male pedicels based on transcriptomic analysis (Related to Figure 6D).**

**Supplementary file 3: Interactive network of enrichment GO analysis of upregulated genes in male pedicels compared to female pedicels based on proteomic analysis (Related to Figure 6H).**

**Supplementary file 4: Interactive network of enrichment GO analysis of upregulated genes in female pedicels compared to male pedicels based on proteomic analysis (Related to Figure 6H).**

**Supplementary file 5: Interactive network of enrichment GO analysis of upregulated genes in male pedicels compared to male heads based on transcriptomic analysis (Related to Supplementary Figure 4A).**

**Supplementary file 6: Interactive network of enrichment GO analysis of upregulated genes in male heads compared to male pedicels based on transcriptomic analysis (Related to Supplementary Figure 4A).**

**Supplementary file 7: Interactive network of enrichment GO analysis of upregulated genes in male pedicels compared to female heads based on transcriptomic analysis (Related to Supplementary Figure 4B).**

**Supplementary file 8: Interactive network of enrichment GO analysis of upregulated genes in female heads compared to male pedicels based on transcriptomic analysis (Related to Supplementary Figure 4B).**

**Supplementary file 9: Interactive network of enrichment GO analysis of upregulated genes in male pedicels compared to male heads based on proteomic analysis (Related to Supplementary Figure 4C).**

**Supplementary file 10: Interactive network of enrichment GO analysis of upregulated genes in male heads compared to male pedicels based on proteomic analysis (Related to Supplementary Figure 4C).**

**Supplementary file 11: Interactive network of enrichment GO analysis of upregulated**

**genes in male pedicels compared to female heads based on proteomic analysis (Related to Supplementary Figure 4D).**

**Supplementary file 12: Interactive network of enrichment GO analysis of upregulated genes in female heads compared to male pedicels based on proteomic analysis (Related to Supplementary Figure 4D).**

**Supplementary Table 1**: **Gene ontology enrichment analysis comparing male pedicels with male or female heads.**

Enriched Gene Ontology (GO) terms (biological processes) and annotated upregulated and downregulated genes (adjusted p value < 0.1) in male pedicels compared to male or female heads.

